# Cryptic binding sites are detected but not ranked: coverage, conversion, and the limits of detector consensus

**DOI:** 10.64898/2026.08.11.743381

**Authors:** Clayton W. Moore

## Abstract

Cryptic-pocket prediction is compared almost entirely by top-*n* recovery, which conflates two separable abilities: proposing a candidate at the true site, and ranking it highly enough to be seen. We retained per-candidate overlaps for five candidate-generation methods, evaluated in six configurations, on the CryptoBench test fold. Coverage spanned 14.5 points; conversion of coverage into top-five recovery spanned 36.7. Two rankers over an identical candidate set differed by 10.6 recovery points at equal coverage, isolating ranking exactly. Union coverage saturated at 92.2%, reaching 98.6% on sites of at least eight residues, with residual failures concentrated on small sites. Injecting synthetic competitors drawn from each target’s own wrong-answer score distribution, holding the true site, candidate pool and ranker fixed, reduced top-five recovery by 16.8 points on training folds and 17.0 points on the held-out test fold. Pooling detectors consequently gains nothing at a budget of five and 11.8 points at twenty. We also report two defects in the evaluation apparatus itself, each of the kind this paper is about. Matching an interface site’s hyphen-joined chain set against single-chain residue labels by string equality silently removed all 38 interface structures of the fold from every method’s row at once. Scoring residue overlap without the chain credited a top-five candidate sharing no residue with the annotated site, its center 36.4 A away. Both are fixed; neither moves a number here, since this cohort is single chain, where the two metrics are provably identical. On the repaired full fold the one configuration re-measured on it, Lacuna 1.5.0 under its default ranker, widens the gap between finding a site and ranking it rather than closing it: 18.0 points across all 222 structures against 15.1 on the 179 analyzed here.

---

A cryptic binding site is one that is not apparent in a protein’s unbound structure and opens only on ligand binding or thermal fluctuation [1,2]. Because the apo structure frequently contains no open cavity at the site, cryptic sites are the hardest case for binding site prediction and among the most valuable, since they expand the set of proteins considered druggable [3].

Progress is measured almost entirely by top-*n* recovery: the fraction of targets for which a qualifying pocket appears within a method’s first *n* proposals, usually with *n* = 1, 3, or 5 [4,5]. This single number combines two abilities that have no necessary relationship. A method must first propose a candidate that overlaps the true site, and must then rank that candidate above its own alternatives. A method that proposes well and ranks poorly is indistinguishable, under top-*n*, from one that never proposes the site at all.

That conflation matters because the field it summarizes is architecturally diverse. Binding site prediction now spans geometric cavity detection [6], machine-learned scoring of surface points [7], 3D convolutional and surface-based deep learning [8,9,10,11], graph neural networks [12,13], probe-based energetic mapping [14], tree-ensemble methods [15], and scoring built on learned protein representations [16,17,18]. Cryptic sites in particular have driven a parallel line of work on conformational sampling, from normal mode analysis [19] and pocket detection over molecular dynamics [20] to generative ensemble models [21,22,23,24]. These approaches differ in what they propose and in how they order what they propose, and top-*n* prices both at once.

The reason the split has not been measured is mundane: it requires keeping information every pipeline discards. Benchmarks record whether a hit occurred within the top five and throw away everything below. Recovering coverage after the fact is impossible without re-running every method.

An earlier version of this study retained per-candidate overlaps for four detectors and reported the decomposition [25]. Three questions it raised could not be answered with those data, and this study is built to answer them. Whether the decomposition is peculiar to the four methods chosen, which we address by adding two more and by separating one method’s two rankers. Whether the detection gap would close if more architectures were added, which requires a union computed over enough methods to see whether it saturates. And whether candidate burden merely correlates with poor ranking or causes it, which requires an experiment rather than an observation.

Answering the third question also required correcting the first version. It reported that added coverage converts to recovery at about 85% below roughly fifteen candidates and about 51% above, implying a threshold. An independent breakpoint search over this larger dataset does not reproduce that threshold: the best descriptive split falls at seven candidates, 23.6% of bootstrap resamples select an edge of the search grid, and a continuous log-linear model fits better than any step (AIC 2342.1 against 2364.0). The competition effect is real and, as shown below, causal. Its functional form is not a threshold, and we no longer claim one.

This version adds a finding of the same character about the apparatus rather than the methods. Auditing the harness that produced the candidate collection analyzed here turned up two defects, one in how the benchmark’s own chain encoding was read and one in how residue overlap was scored. Both were silent, both change what a comparison means, and neither moves a number in this paper, for reasons we give and verify. They are reported because they generalize past the harness that carried them.

## Results

### Coverage varies little between detectors; conversion varies enormously

Table 1 gives the decomposition on the designated test fold. Coverage spans 14.5 points across the six configurations and conversion spans 36.7.

**Table 1.** Coverage and conversion by configuration. Designated CryptoBench test fold, n = 179.

| CONFIGURATION | Candidates | Coverage | Top-5 | Conversion |
|---|---|---|---|---|
| fpocket | 18.9 | 73.7% | 43.6% | 59.1% |
| P2Rank | 6.7 | 65.9% | 63.1% | 95.8% |
| IF-SitePred | 23.3 | 70.9% | 62.0% | 87.4% |
| MDpocket | 13.6 | 59.2% | 44.1% | 74.5% |
| Lacuna | 20.2 | 71.5% | 53.6% | 75.0% |
| Lacuna-PLM | 20.2 | 71.5% | 64.2% | 89.8% |

The ordering reverses depending on which ability is measured. fpocket has the highest coverage of any method tested and the lowest top-five recovery. P2Rank proposes the fewest candidates of any single-structure method and converts almost all of them. Under top-five alone these two methods are separated by 19.5 points and the reason is invisible.

**Figure 1.**
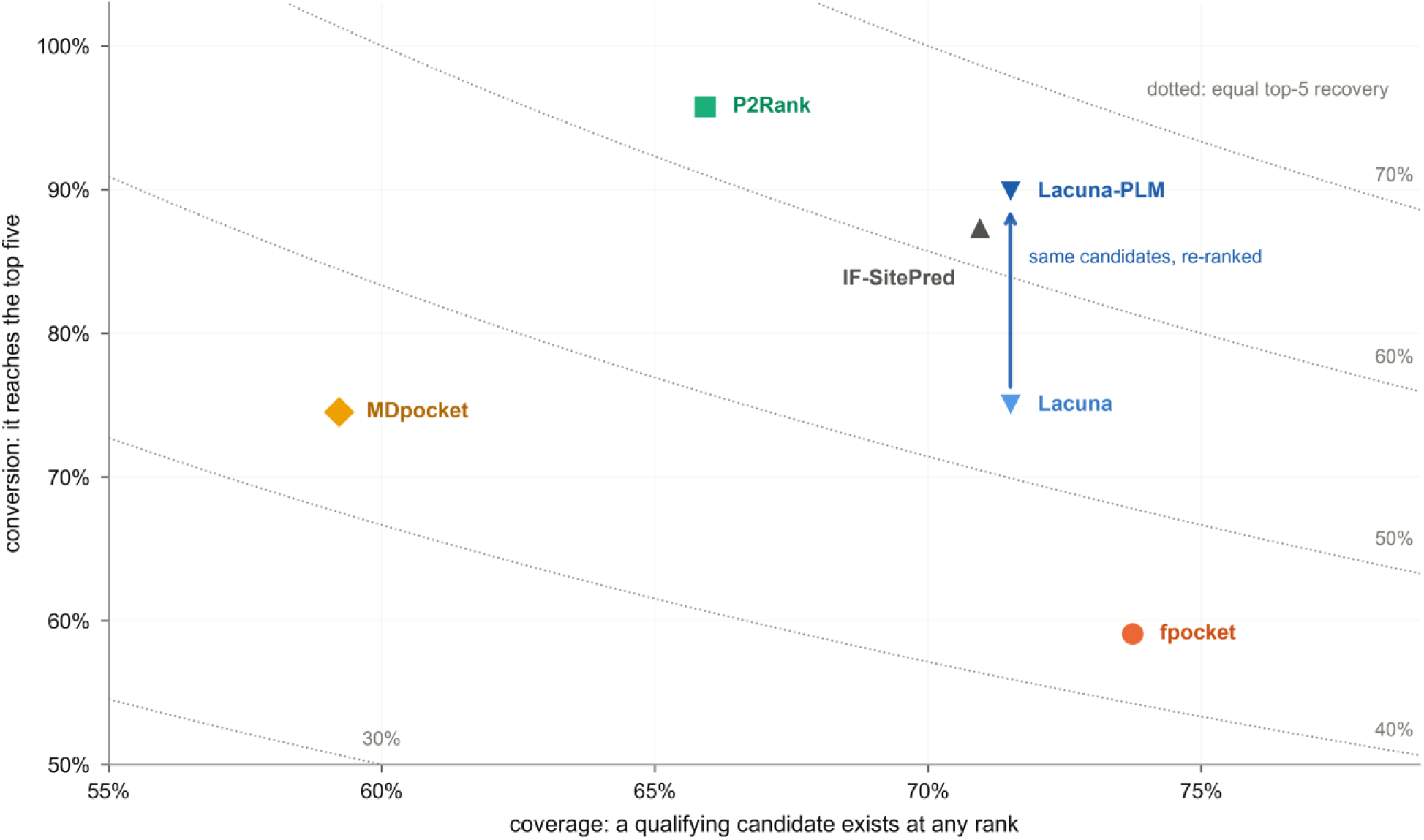
Recovery is the product of two separable abilities. Top-five recovery equals coverage multiplied by conversion, so equal-recovery contours are hyperbolas (dotted). Six configurations on the designated CryptoBench test fold, n = 179. fpocket and P2Rank are separated along the diagonal of the plane: fpocket has the widest coverage in the set and the lowest conversion, while P2Rank sits mid-range in coverage and converts almost everything it proposes. Their marks lie two equal-recovery contours apart, 43.6% against 63.1%, and a bar chart of recovery alone shows that 19.5 point difference as a single number and hides its cause. The pair that does land on nearly onecontour is fpocket and MDpocket, 43.6% and 44.1%, which arrive there from the widest coverage gap in the set, 73.7% against 59.2%: equal recovery is not equal ability. The vertical arrow joins Lacuna’s two rankers, which operate on an identical candidate set: coverage is fixed by construction and only conversion moves.

### The same candidates, ranked differently

The cleanest evidence that the two abilities are separable does not require comparing methods at all. Lacuna’s two rankers operate on an identical candidate set: the same detector, the same ensemble, the same clustering, differing only in how the resulting sites are ordered.

Their coverage is identical at 71.5%, necessarily, because the candidates are the same. Their top-five recovery differs by 10.6 points, 53.6% against 64.2%. A within-method comparison cannot be attributed to differing candidate semantics, differing pocket definitions, or differing numbers of proposals, all of which complicate cross-method comparison. It isolates ranking exactly.

**Figure 2.**
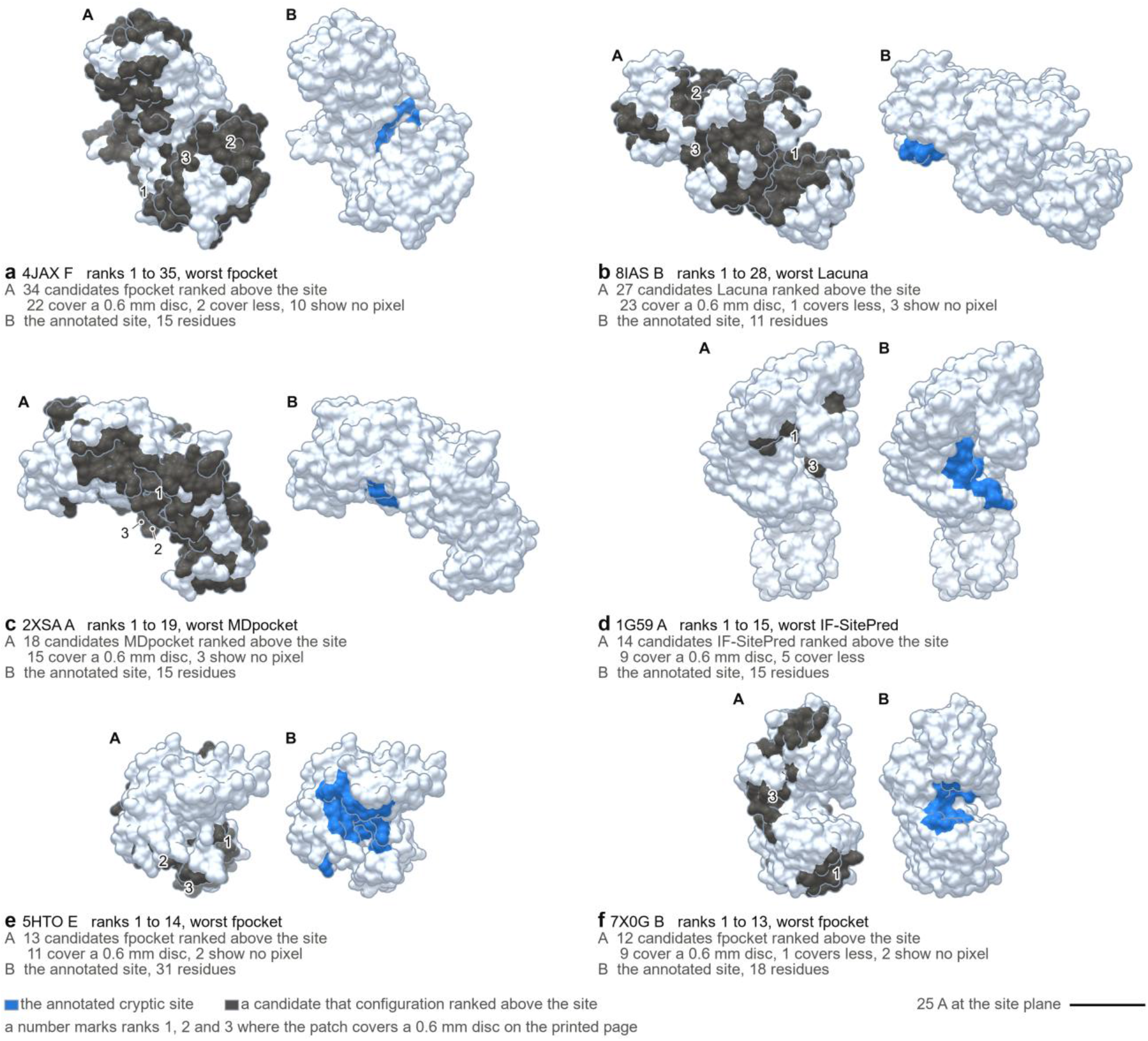
The same site, ranked very differently across configurations. Six test-fold targets for which every configuration proposes a qualifying candidate, ordered by how far apart they rank it. Each target is a pair of frames on one camera and one crop, so the two differ only in what is colored: A draws every candidate the configuration that ranked the site worst put above it, with the frame’s own line naming how many that is, and B draws the annotated cryptic site alone, in blue. Every candidate patch is painted in the one flat grey, at every rank. That grey means only that the configuration ranked the candidate above the site: it does not carry rank, and the variation of tone within a patch is the renderer’s lighting on the molecular surface, which spreads a single ink over twenty or more lightness steps and carries no information. Rank is carried instead by the drawn numbers and by each frame’s own count. Ranks 1, 2 and 3 carry a number wherever that patch covers at least the area of a disc 0.6 mm across on the printed page, either on the patch or on a short diagonal leader ending in a dot on it. Nothing is drawn where no record exists: a patch is the residues that configuration stored for that candidate and no more, less any it shares with the site, which only B draws. Each frame’s own line gives how many of its candidates cover that 0.6 mm disc of printed page, then names the two remainders wherever either exists: those that cover less than it, which are drawn but as specks, and those that put no pixel on the page at all. Twenty candidates across the six panels fall in that last class, and eighteen of them register nothing in a render of their own surface either, so they have no surface visible from that camera; the other two are slivers the printed silhouette outline covers. Those counts are measured on the printed render itself, not on the geometry behind it. Panel captions give the range across all six and name the worst. In (a) fpocket returns thirty-four candidates before the site another method places first. Panels share a common scale and a common orientation procedure, and because the camera is perspective the bar is 25 A at the site plane.

### Adding detectors saturates

If the field’s remaining failures were detection failures, adding architectures should keep expanding the union. It does not. Adding configurations greedily, each step taking whichever contributes most new coverage, gives 73.7% for fpocket alone, 89.9% once IF-SitePred joins it, 91.6% with P2Rank and 92.2% with Lacuna, after which nothing further is gained (Supplementary Table 1).

The fifth architecture to be added, MDpocket, contributes nothing measurable, and it is the one most structurally unlike the others, accumulating a pocket-frequency grid over an ensemble rather than scoring a single structure. Lacuna-PLM contributes nothing either, but that zero is a tautology rather than a result: it shares Lacuna’s candidate set exactly and therefore cannot change coverage at all. The union is saturated at roughly 92%, and the 7.8% that remains is not a matter of having tried too few methods.

**Figure 3.**
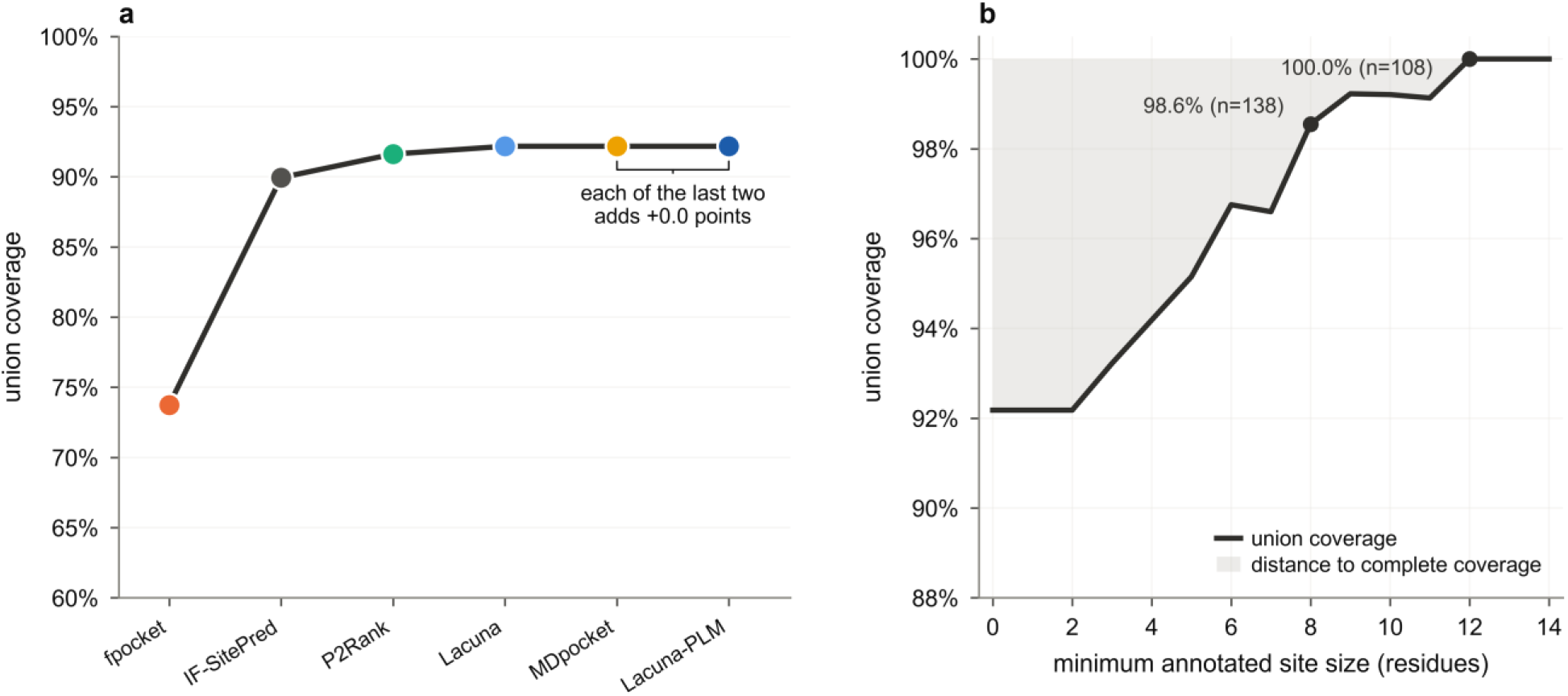
The union saturates, and the remaining gap is concentrated in small sites. (a) Configurations added greedily, each step taking whichever adds most coverage. Two methods reach 89.9% and four candidate generators reach 92.2%; the fifth architecture adds nothing measurable, and the sixth marker is a second ranker over an already-counted candidate set, which cannot add coverage. (b) Union coverage against a floor on annotated site size. Shading marks the distance to complete coverage. Targets no method proposes have a median site of 5 residues against 15 for the rest, and the union reaches 98.6% once sites of fewer than eight residues are excluded.

### What the remaining 7.8% is

The targets no method proposes are not a random sample. Their annotated sites have a median of 5 residues against 15 for the rest of the fold, and 92.9% of them are eight residues or smaller against 22.4% of covered targets. Restricting to sites large enough that the criterion has room to operate raises union coverage from 92.2% (n = 179) to 96.8% at a floor of six residues (n = 154), 98.6% at eight (n = 138) and 99.2% at ten (n = 126), while the best single configuration’s top-five rises from 64.2% to 83.3% over the same range (Supplementary Table 2).

This is not a scoring artifact, and we tested it as one. Every target in the fold had at least one candidate whose size could in principle reach the threshold given perfect overlap, so no target was unscoreable by construction. A size-independent centroid criterion rescues only one of the fourteen invisible targets. Small cryptic sites are genuinely harder to detect, not artifactually penalized.

The consequence for the field’s arithmetic is nonetheless substantial, and it is worth splitting rather than quoting whole. On sites of at least eight residues the six configurations between them propose a qualifying candidate for 98.6% of targets, while the best single configuration returns one within five for 79.7%, a gap of 18.8 points. That gap is two different problems. Lacuna-PLM, the best configuration at this floor, covers 87.0% on its own. So 7.2 points are sites it proposes and then ranks outside five, which better ranking alone would recover, and 11.6 points are sites it never proposes at all, which only pooling with another method can reach. Neither half requires a new detection architecture, but they are not addressable by the same means, and reporting the 18.8 as a single ranking deficit would overstate what a better ranker can do.

**Figure 4.**
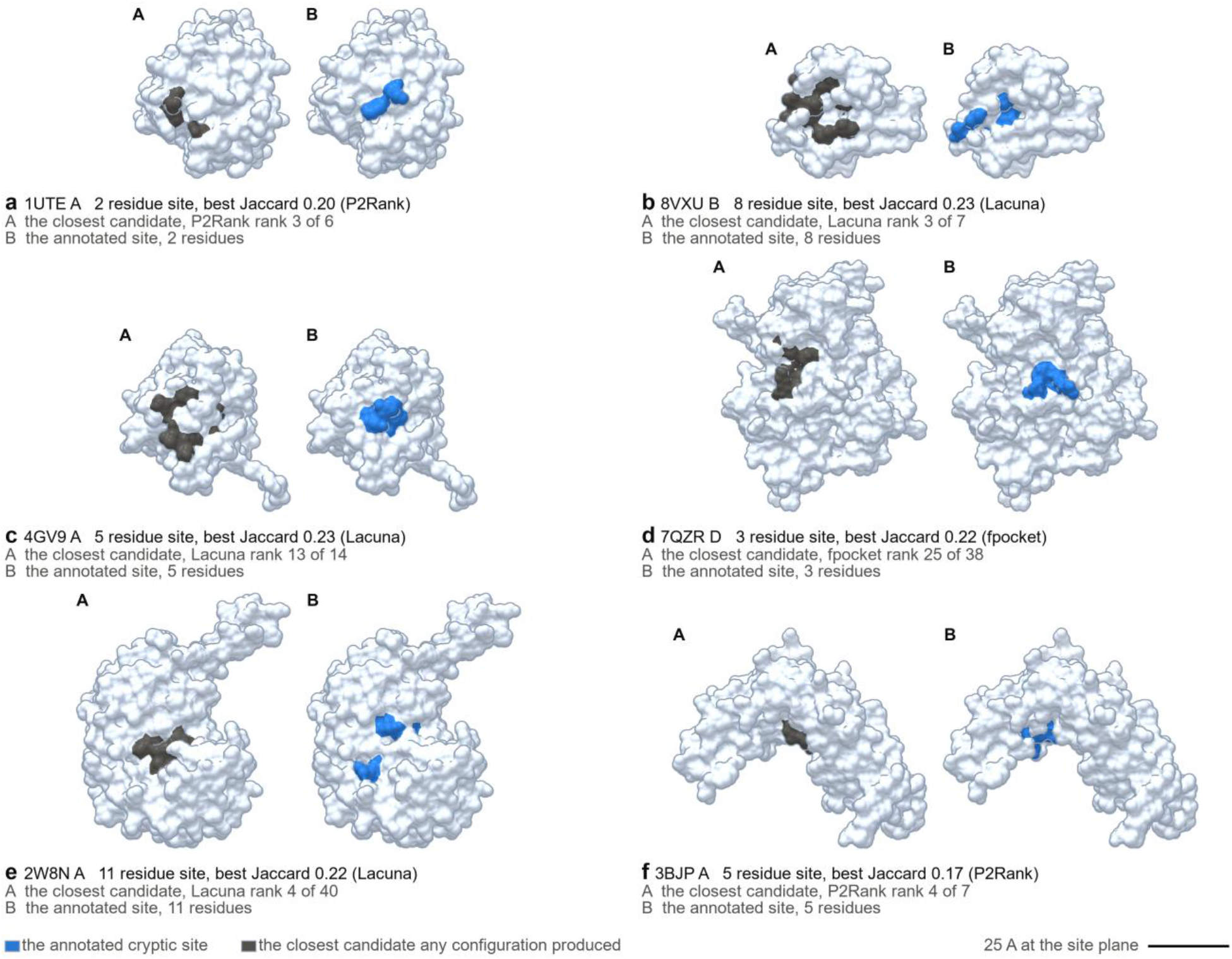
What the irreducible remainder looks like. Six of the fourteen test-fold targets for which no configuration proposes a qualifying candidate. Each target is a pair of frames on one camera and one crop, so the two differ only in what is colored: A is the closest candidate any configuration produced, in the same flat candidate grey Figure 2 uses, and B the annotated site alone, in blue. Nothing is drawn where no record exists: a patch is the residues stored for it and no more, less any it shares with the site, which only B draws. These are near misses rather than wild ones: the best Jaccard of all six shown runs from 0.17 to 0.23 as each panel prints it, short of the 0.25 threshold, and in every target the closest candidate abuts or overlaps the site. The failures concentrate on small annotations, with a median of 5 residues against 15 elsewhere in the fold. They are nonetheless genuine detection failures rather than scoring artifacts: every target in the fold had candidates whose size could in principle reach the threshold, and a size-independent centroid criterion recovers only one of the fourteen. Panels share a common scale, and because the camera is perspective the bar is 25 A at the site plane.

### Candidate burden causally reduces top-*k* conversion under a fixed ranker

Conversion falls as candidate burden rises. Among covered targets, the rank correlation between candidate count and top-five success is -0.257 pooled and negative within every configuration separately. It survives protein length, which is not significant, and detector fixed effects, which are large.

Correlation is not enough here, because burden is not assigned. Coverage rises with burden almost mechanically, so conditioning on coverage and regressing conversion on burden conditions on a collider: among covered targets, low-burden cases were covered easily while high-burden cases may be covered only because extra candidates caught a marginal site. Two experiments break that.

### Varying the ensemble

Running Lacuna at 1, 3, 5, 10 and 20 conformers changes burden on the same targets with the same detector, ranker and criterion. This sweep runs on the pooled CryptoBench train folds, not on the 179-target test fold used above. Over the 720 train-fold targets common to every level of the sweep, mean burden rises from 6.6 candidates at one conformer to 21.8 at twenty. Restricted to the 330 of those train-fold targets covered at both ends, so coverage selection cannot operate, that growth costs 7.9 points of top-five recovery (95% CI 5.2 to 10.9) and pushes the first qualifying candidate down by 0.78 ranks (95% CI 0.53 to 1.05).

### Injecting competitors

More conformers changes what the candidates are as well as how many. To hold quality exactly fixed, we instead spliced synthetic competitor entries into the ranked list, each scored by a draw from that target’s own non-qualifying score distribution, leaving the real candidates, the annotated site and the ranker untouched. Adding 40 such entries costs 16.8 points at top-five (95% CI 14.0 to 19.7), and every top-*k* from *k* = 3 to *k* = 20 shows an interval excluding zero. Nothing about the target has changed except how many competitors its ranked list contains.

### The same result on held-out data

The ranker used above was fitted on CryptoBench training data, so the experiment as described measures displacement on targets that helped fit the scorer. Repeating it unchanged on the designated test fold, which the ranker never saw, gives 17.0 points at top-five (95% CI 11.3 to 23.0) on the 130 eligible targets. The point estimate is within 0.2 points of the train-fold value and every top-*k* from 3 to 20 again excludes zero (Figure 5c). The effect is not an artifact of evaluating a model on its own training data.

**Figure 5.**
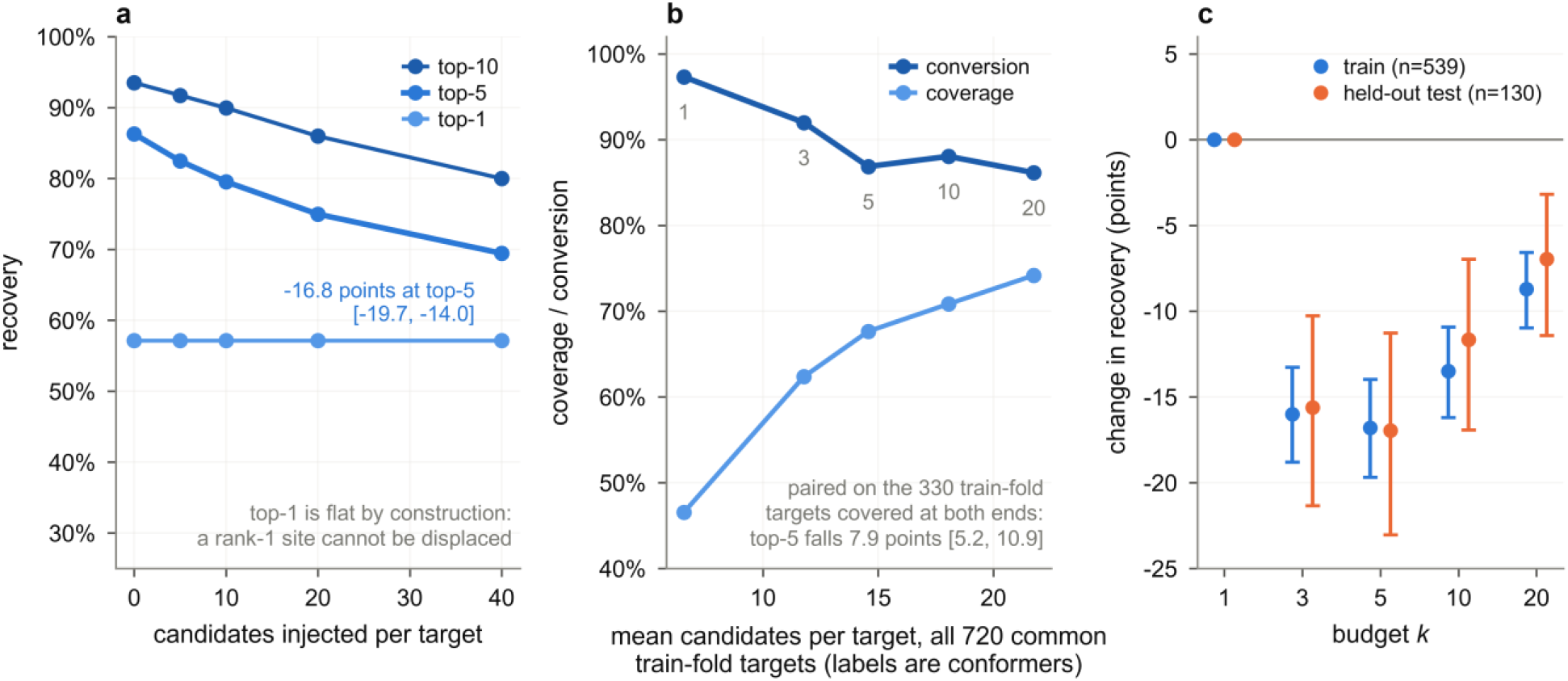
Burden displaces findable sites, with quality held fixed. (a) Recovery against the number of synthetic competitor entries spliced into the ranked list, each scored by a draw from that target’s own non-qualifying score distribution, so that candidate quality, the true site and the ranker are all unchanged and only the number of competitors moves (n = 539 train-fold targets, 20 draws per budget). Top-five falls from 86.3% to 69.4%. Top-one cannot move under this design and does not, for the reason given in the text. (b) The same effect reached a different way, by growing the conformational ensemble from 1 to 20 conformers: coverage rises at every step while conversion ends 11.2 points lower, falling at every step except 5 to 10 conformers, where it recovers 1.2 points. Restricting to the 330 train-fold targets covered at both ends removes coverage selection as an explanation. (c) The injection experiment on the pooled train folds against the identical procedure on the held-out test fold, which the ranker never saw. Paired change in recovery at the largest budget, with 95% bootstrap intervals. The two agree at every *k*, so the effect does not depend on evaluating the ranker on its own training data.

The design cannot move top-one, and does not: the paired effect at *k* = 1 is exactly zero, with a bootstrap interval of [0, 0]. That null is structural rather than evidential. Injected scores are drawn from the target’s own non-qualifying candidates, so the largest score any injection can carry is the largest wrong score the target already produced, and a site ranked first already outscores it. No number of draws can displace it. The flat line in Figure 5a is therefore the price of holding quality exactly fixed, not evidence that burden stops mattering at the tightest budget.

Injecting candidates drawn from *other* proteins does displace the true site from rank one, in 28.4% of cases. The ranker is fitted on within-structure pairs and nothing constrains its scores to be comparable across structures, so that variant measures score miscalibration on top of burden. We report the within-target version as the causal estimate for exactly that reason, and return to the cross-structure result below, where it bears on pooling.

### Consensus pays only outside the conventional budget

Because burden costs ranking, pooling configurations is not free. The pooled analysis is computed on the earlier four-detector candidate collection, fpocket, P2Rank, IF-SitePred and Lacuna under its PLM-assisted ranker, on the 178 targets all four ran on, rather than on the six configurations of Table 1; Methods gives the reason and the size of the difference. Spending a fixed budget across those four detectors rather than within the best single one gains nothing at a budget of five (66.3% against 66.3%), 7.9 points at ten (77.0% against 69.1%) and 11.8 points at twenty (83.7% against 71.9%) (Supplementary Table 3).

At the conventional budget of five the paired difference between the allocation fitted on the train folds and the best single detector is +0.0 points (95% CI -2.8 to +3.4); for the round-robin pooling of the table it is also +0.0 points, with a wider interval (95% CI -5.6 to +5.6). Consensus is worth nothing there on either strategy. By twenty the single-configuration curve has saturated at its own coverage ceiling while the pooled curve continues toward the union.

Recomputing the same round-robin pooling over the six configurations of Table 1, on this paper’s 179 targets, preserves the shape and not the magnitudes: -2.8 points at a budget of five (95% CI -8.9 to +3.4), +3.9 at ten (95% CI -1.7 to +9.5) and +10.1 at twenty (95% CI +5.0 to +15.6). Round-robin over six configurations spends two of its slots on one candidate set, because Lacuna’s two rankers share candidates exactly, which is why it falls behind the best single configuration at the tightest budget. The conclusion survives either accounting.

The field reports top-five. On CryptoBench, under the pooling strategies tested here, that is precisely the regime in which detector consensus provides no measurable gain.

### Two defects in the evaluation apparatus, and what each does to a comparison

The harness that produced the candidate collection analyzed here carried two defects. Both were found after version 2 of this study was posted, both are now fixed, and neither changes any number reported above, which we verify below. We report them here rather than as an erratum, and in Results rather than in Discussion, because they are measurements of exactly the kind the rest of this paper makes: properties of evaluation practice that are invisible in a published summary and that change what a comparison means. Neither is specific to the tool whose harness carried them.

### A benchmark’s chain encoding can delete a structural class

CryptoBench names an interface site by the hyphen-joined set of chains it spans, “M-O-P”, “G-H”, “AAA-CCC”, “B-D-F-H”, while each annotated residue of that site carries a single chain id drawn from the set. The harness compared each residue’s chain prefix against the joined string for equality. For a multi-chain site that equality never holds, so the residue set came back empty and the entry was dropped: counted as an anonymous skip, with no structure named and no reason recorded. Thirty-eight of the 222 entries of the designated test fold are interface sites encoded this way, and all 38 were removed, from every method’s row at once, since the collection script takes its site definition from the same function (benchmarks/collect_candidates.py imports main_pocket and the residue ceiling from benchmarks/cryptobench_benchmark.py).

What that does to a published comparison is not noise. The loss is perfectly paired, so it does not obviously tilt a ranking, and that is what makes it hard to notice: every row shrinks by the same structures, the table still balances, and the only visible trace is a denominator smaller than the benchmark’s. What disappears is a structural class. Interface sites, all 38 of them homomultimers in this fold, are simply not in the measurement, so no conclusion in a paper scored this way is supported for them, including conclusions the paper does not know it is making about them.

### Residue overlap without the chain credits the wrong chain

The residue Jaccard used to score a candidate reduced pocket lining labels, which carry a chain, to bare sequence numbers. On a single-chain structure this is harmless. On a homomultimer, where every chain repeats the same numbering, a candidate lining one chain can be credited against annotated residues on another. The clearest measured case is 3x0x, whose site spans chains E and G: the candidate at rank 3, 61 residues of lining, shares no residue at all with the annotated site and its center lies 36.4 A from the site’s C-alpha centroid, yet it scored a residue Jaccard of 0.29 chain blind, above the 0.25 threshold used here, against 0.00 chain aware. Its chain-blind recall is 0.95, 18 of the site’s 19 residue numbers matched and every one of them on the wrong chain. Reading that record also requires care of the kind this subsection is about: the stored residue Jaccard is a maximum over the rank block, while the rank and centroid distance stored beside it belong to the block’s nearest-centroid candidate, so the three fields need not describe one candidate and on 3x0x they do not. Scoring the chain also repairs the annotation itself, since the chain-blind site set collapses repeated numbers and was short by one to three residues on 8 of the 38.

This defect is worse for comparison than the first, because it is not uniform across methods. The credit goes to whichever method happens to propose a pocket on the other chain of a homomultimer, and methods differ in how many candidates they propose and where. In the one comparison in which both arms could be put on the corrected metric, Lacuna’s fitted ranker against the analytic rule it replaced, the chain-blind metric inflated the weaker arm more: it was carrying 11.1% of the analytic arm’s hits against 5.3% of the fitted arm’s, so the measured gap between the two grew by about half a point, from 27.5 points to 27.9, once the chain was kept. Only the fitted arm could be re-scored from stored per-structure records; the analytic arm’s stored file predates the fix and keeps no per-cluster lining residues, so that arm was re-run on the full fold with the corrected harness. On the full fold the correction costs 7 of the 38 interface structures their top-five hit, and the full-fold top-five figure for that tool falls from 133/222 to 126/222, 59.9% to 56.8%. No single-chain structure moves by any amount.

### Why neither defect moves a number in this paper

The cohort analyzed here is 179 single-chain structures (Methods), so the first defect determines which structures are present and is accounted for there, and the second cannot reach them: on a structure with one chain, scoring residue overlap with the chain and without it are the same operation. That identity is verified rather than assumed. Over all 184 single-chain entries of the fold, in every rank block and every field, under both ranking arms, the chain-aware and chain-blind residue Jaccard and recall agree exactly, 0 mismatches in 2,208 comparisons; this paper’s 179 are a subset of those 184. The provenance of that check, the fix and every figure quoted above is recorded in benchmarks/PROVENANCE_cryptobench_full_fold_2026-11-16.md.

### The gap is wider on the repaired full fold

With both defects fixed the whole 222-structure fold can be scored, and one configuration has been re-measured on it: Lacuna 1.5.0 under its default ranker, chain-aware metric, all 222 attempted and all 222 scored with nothing skipped. Coverage at any rank is 166/222 (74.8%) and top-five recovery 126/222 (56.8%), a conversion of 75.9% and a gap between finding a site and ranking it of 18.0 points. The same release on this paper’s 179-structure cohort gives coverage 130/179 (72.6%), top-five 103/179 (57.5%), conversion 79.2% and a gap of 15.1 points. Restoring the 43 structures this paper’s cohort leaves out therefore widens the separation this paper is about rather than closing it, on a cohort 24% larger.

The version is stated because it differs from the rest of the paper: every Lacuna row in Table 1 is version 1.0.1, and this measurement is 1.5.0 (Methods). For that one configuration the 38 interface structures look ordinary at top five, 57.9% against 56.5% for the 184 single-chain structures, but decomposed in this paper’s own currency they are not. Coverage on the interface cohort is 84.2% (32/38) against 72.8% (134/184), while conversion of that coverage into top-five recovery is 68.8% (22/32) against 77.6% (104/134): easier to propose, harder to rank, by 11.4 points and 8.9 points respectively. The two effects very nearly cancel in the top-five number, which is the paper’s own argument reappearing in the cohort that was missing from the measurement. The gap between finding a site and ranking it is 26.3 points on the interface structures against 16.3 on the single-chain ones, and they are what lifts the full-fold gap to 18.0. The power caveat is as important as the decomposition: n = 38 cannot resolve either difference on its own, so this is the shape of the effect in its purest observed form and not an independent measurement of its size. It is also one method and one version and it is not evidence about the other four.

## Discussion

Five practical conclusions follow.

### Report coverage and conversion separately

They are separable abilities, they vary independently across methods, and a single top-*n* number obscures both. A method with 74% coverage and 59% conversion needs different work from one with 66% coverage and 96% conversion, and top-*n* cannot distinguish them. The per-candidate data required is already computed by every method and merely discarded.

### Detection is close to saturated among the methods tested

Two methods reach 89.9% coverage and four candidate generators reach 92.2%, after which a fifth and deliberately dissimilar architecture adds nothing measurable. On sites of at least eight residues the union reaches 98.6%. Adding increasingly distinct architectures therefore produced sharply diminishing returns on CryptoBench; we cannot exclude that some future method proposes what none of these do, but the returns observed here fall to zero well before the benchmark is exhausted. Effort spent proposing more candidates is, on this evidence, worth very little compared to effort spent ranking existing ones better, and the remaining detection failures concentrate on small sites rather than being spread across the benchmark.

### Candidate burden is a cost, not merely a correlate

Two experiments, one varying the ensemble and one injecting candidates of fixed quality, both show that adding candidates pushes findable sites out of a fixed budget. This reframes a class of interventions: any method that improves coverage by proposing more must pay for it in ranking, and the payment is measurable.

### Consensus needs a budget to work in

Pooling detectors is worth nothing at five candidates and 11.8 points at twenty. A field that reports top-five will conclude that consensus does not help, and will be right about top-five and wrong about consensus.

### Audit the harness, not only the method

The two defects reported above had the signature this paper is about: each was invisible in the summary number and each changed what a comparison meant. One deleted a structural class from every row at once, leaving a comparison that looked perfectly paired and was quietly about less of the benchmark than it claimed. The other credited a candidate on one chain of a homomultimer against annotated residues on another, which inflates whichever method proposes there and so is not uniform across methods. Neither belongs to the tool that carried them. Any evaluation that matches an annotation’s chain field by string equality inherits the first, and any method scored by residue-number overlap on a structure with more than one chain inherits the second. Two practices would have exposed both: report per structure what was scored and what was skipped and why, with skips counted as misses in a stated strict rate, so that a silent drop cannot survive a run; and compute residue overlap on (chain, number) pairs wherever the structure has more than one chain, recording the chain-blind value alongside so that the size of the difference stays auditable.

### Several limitations remain

The field-level comparisons rest on a single benchmark, CryptoBench, and the five candidate generators, although architecturally diverse, do not exhaust the published binding-site prediction literature. Whether the magnitude of the coverage-conversion separation and of union saturation generalizes to other cryptic-site benchmarks remains untested. The cross-detector comparison covers 179 single-chain structures of a 222-structure fold, for the reasons set out in Methods, so nothing here measures how any of these methods behaves on an interface site; the one configuration re-measured on the full fold finds those sites easier to propose and harder to rank, at an n that cannot resolve either difference, and that is one method. All Lacuna rows are version 1.0.1, and a later release of the same tool differs by 3.9 points of top-five recovery on this cohort, which is larger than several effects reported here; the comparison is held at one version on purpose, because the other four methods cannot be re-run under matched conditions. The candidate-burden experiments are Lacuna-only, because they require re-running a detector under controlled conditions; whether the effect size transfers to other architectures is untested. The ensemble-sweep experiment uses the pooled train folds and has no held-out counterpart, since varying the conformer count requires re-running the detector rather than re-reading a stored list; only the injection experiment is replicated on the test fold. The injection design holds candidate quality fixed at the cost of being unable to perturb rank one at all, so at the tightest budgets it gives a lower bound on burden rather than an estimate. It also manipulates the ranked list rather than protein structure, so it isolates ranking competition specifically and says nothing about whether real additional pockets would compete more or less strongly than the synthetic entries do. The pooled consensus reported here allocates a budget across detectors but does not re-rank their candidates jointly, which would require calibrating scores across methods. That calibration is not straightforward: the 28.4% displacement rate under foreign injection shows the ranker’s scores carry no cross-structure meaning, which is a prerequisite for pooling. And the designated test fold is not fully blind, since the tool developed here was measured against it during development; the comparator detectors were run at published defaults and were not.

## Methods

### Benchmark

CryptoBench [4] supplies apo structures with cryptic site residues annotated from a holo counterpart, split into homology separated folds. We report the designated test fold (n = 179) as the primary result and the pooled train folds (n = 743) where a larger sample aids resolution.

### Cohort

The designated test fold has 222 entries and this study analyzes 179 of them. The arithmetic is as follows. Thirty-eight entries are interface sites whose chain field is a hyphen-joined set, and the collection harness dropped every one of them through the chain-matching defect reported in Results. Four more exceeded a 700-residue ceiling in the same harness and were skipped by it: 2aka A, 4zoe B, 7f4y B and 7o1i A. One further structure, 2v6mD, produced no output row at all from two of the six configurations, IF-SitePred and MDpocket, and so cannot enter a paired comparison. 222 minus 38 minus 4 minus 1 is 179.

Both harness defects were fixed on 2026-11-16, after version 2 of this study was posted; the ceiling now stands at 1,200 residues, which covers the whole fold, and every unscored structure is named with a reason. The exclusion of the 38 interface entries is therefore no longer a silent loss but a stated choice, and the reason is that the comparison cannot be re-made correctly today. P2Rank is not installed in this environment and no record states which build produced the stored rows. IF-SitePred’s vendored code cannot score a multi-chain site: it takes the first chain and exits successfully. The stored Lacuna, Lacuna-PLM and MDpocket rows share a single conformational ensemble generated by version 1.0.1, whose sampler was float32 and thread dependent and so cannot be reproduced. Scoring 38 homomultimers under a metric that can credit the wrong chain would be worse than leaving them out, and scoring them under the corrected metric would require re-running all six configurations, which is not possible here. The full fold is reported for the one configuration that could be re-measured (Results).

### Zero-candidate outcomes

A target enters the analysis only when every detector produced output for it, so all comparisons are paired. Producing no candidates counts as output: it is a prediction that the structure has no detectable site, and it is scored as a miss rather than dropped. This is the only reason the test fold here numbers 179 against 178 in the earlier version [25], whose loader discarded rows with empty candidate lists. The single added target, 1bk2A, is one for which five of the six configurations return nothing at all.

The choice is not cosmetic. Six targets yield no IF-SitePred candidates, and scoring them as misses rather than dropping them moves its coverage from 73.4% to 70.9%. A benchmark that silently drops empty predictions therefore reports up to 2.5 points of coverage that the method did not earn; the corresponding inflation is 0.7 points for P2Rank and 0.3 to 0.4 points for the rest. We recommend that empty predictions be reported and scored rather than filtered.

### Detectors

Five candidate-generation methods spanning the architectural range and seventeen years, evaluated in six configurations: fpocket 4.2 [6], purely geometric, alpha spheres on a Voronoi tessellation; P2Rank 2.5.1 [7], a random forest over physicochemical descriptors of solvent accessible surface points; IF-SitePred [16], which scores residues with a LightGBM model over ESM-IF1 [17] inverse-folding embeddings and clusters probe points around the positives; MDpocket [20], which accumulates a pocket frequency grid over a conformational ensemble; and Lacuna 1.0.1 [26,27] under two rankers, one a linear model over geometric and ensemble-derived features and the other adding four features derived from ESM-2 650M [18].

Lacuna’s two rankers share a candidate set exactly and differ only in ordering, so they are two configurations of one generator rather than two detectors. Where a quantity depends only on which candidates exist, as coverage and the union do, the two are interchangeable; we count generators rather than configurations in those places, and say which we are doing.

### A version note

Every Lacuna row here is version 1.0.1, the release current when the collection was made, and the comparison is deliberately held there because the four comparators cannot be re-run under matched conditions. The current release is 1.5.0 [26,28]. On this paper’s 179 structures, under the same Jaccard criterion, its default ranker recovers the site within five for 103 structures (57.5%) against 96 (53.6%) for 1.0.1, with coverage 130 (72.6%) against 128 (71.5%). That 3.9-point difference is larger than several effects reported here, and it is not a clean measurement of a version: the 1.5.0 run was made on Windows and the collection analyzed here in WSL, a platform difference worth +1.7 points of top five in the comparison below, and 1.5.0 also replaced the sampler and changed the rule that decides which residues line a pocket. It is quoted as the size of a confound, not as a measured improvement.

### A shared ensemble

MDpocket and Lacuna both consume a conformational ensemble, generated by anisotropic network model normal mode analysis [19]. The ensemble is generated once per structure and handed to both, so the difference between them isolates detection, aggregation and ranking from conformational sampling. In the earlier version this property was asserted; here it holds by construction.

### One platform

Every detector was run in a single environment. This matters more than it appears: Lacuna 1.0.1’s ensemble derives from an eigendecomposition whose eigenvector signs and near-degenerate mode ordering are decided by the linear algebra backend rather than by the random seed. Running identical code with an identical seed on two platforms left 106 of 180 candidate lists byte-identical, 58.9%; that 180 is this comparison’s own paired cohort, the structures carrying a Lacuna row in both platform dumps, and not the 179 of Table 1. In aggregate the difference is not resolvable (top-five +1.7 points, 95% CI -1.1 to +4.4, n = 180), and results should be reported with the platform stated.

Version 2 of this paper added that per-structure reproduction across platforms is not achievable. That claim is true of the version measured here and is not a property of the approach, and it is dated accordingly: version 1.5.0 of the same tool replaced the sampler, and reports a pocket report identical at any thread count and identical between a Windows and a Linux build on the 40 proteins compared [26,28]. The 58.9% above should be read as a measurement of 1.0.1.

### Criterion

A candidate qualifies if the Jaccard index between its lining residues and the annotated site residues is at least 0.25. Jaccard rather than recall because recall is trivially gamed by proposing larger pockets: ordering candidates by volume alone recovers 77.7% of test-fold structures under a recall threshold of 0.30 but only 52.0% under Jaccard at 0.25, and a trained ranker scores 77.7% under recall as well, so measured that way it is indistinguishable from sorting by size.

### Metrics

For each structure and detector we record the ranked list of per-candidate Jaccards, the lining residues of every candidate, and each candidate’s centroid. Coverage is the fraction of structures where any candidate qualifies at any rank. Top-*k* is the fraction where one qualifies within the first *k*. Conversion is top-five divided by coverage. Confidence intervals are bootstrap percentiles over structures, 20,000 resamples; comparisons between detectors are paired on structure.

Residue overlap is computed on bare sequence numbers in the collection analyzed here and on (chain, number) pairs in the repaired harness. On this cohort the two are not merely expected to agree but measured to: across all 184 single-chain entries of the fold, three rank blocks, two fields and two ranking arms, the two conventions give identical values in 2,208 of 2,208 comparisons, and the 179 analyzed here are a subset of those 184 (Results).

### Candidate injection

This experiment manipulates ranking competition alone, and it does so at the level of the ranked list rather than by generating new pocket geometry. Every real candidate is scored with the production PLM-assisted ranker. Into that scored list we splice *m* additional entries, each assigned a Jaccard of zero, so that it can never qualify and can only displace, and a score drawn with replacement from the empirical distribution of that target’s own non-qualifying candidate scores. We call these synthetic competitor entries rather than candidates, because no pocket is constructed: they are draws from the target’s own wrong-answer score distribution, and are therefore statistically indistinguishable from wrong answers the target already produced. This is the property that makes the manipulation clean. The target’s real candidates, its annotated site and the ranker are all untouched, so candidate quality is held exactly fixed and only the number of competitors moves. The merged list is re-sorted by score and the rank of the first qualifying candidate is re-read.

Eligible targets are those with at least one qualifying candidate and at least two non-qualifying ones, the latter because the procedure needs a within-target null distribution to draw from. Budgets are *m* = 0, 5, 10, 20 and 40, each averaged over 20 independent draws (seed 0; the paired analysis uses seed 1). Intervals are bootstrap percentiles over targets, 20,000 resamples, paired against each target’s own *m* = 0 rank.

We run this twice. The pooled train folds (n = 539) give the powered estimate. Because the PLM-assisted ranker was itself fitted on CryptoBench training data, we repeat the identical procedure on the designated test fold (n = 130 eligible of 179), which the ranker never saw, as a held-out replication.

### Budget allocation

The pooled-budget analysis spends a fixed total of *B* candidates across detectors under three strategies: all *B* within the single detector with the best train-fold top-*B*, round-robin across detectors, and a fixed allocation searched exhaustively on the train folds and reported on the test fold. Hit means any selected candidate qualifies, which needs each candidate’s overlap with the true site and no cross-detector candidate matching.

This analysis runs on the earlier four-detector candidate collection, n = 178 test targets and 734 train, not on the six-configuration collection behind Table 1, and it is the one place in this paper where the two differ. Version 2 described it as pooling across the six configurations; it pools across four detectors, and the description is corrected here. Supplementary Table 3 reports the values, which are unchanged, and its version 2 caption carries the same error. The consequence to be aware of is in the quoted best single configuration. At a budget of five that is Lacuna under the PLM-assisted ranker at 66.3% (118/178), while Table 1 reports 64.2% (115/179) for the same ranker. The 2.1 points decompose exactly: on the 178 structures the two collections share, the earlier Windows collection scores 118 and the single-platform WSL collection 115, seven structures differing and splitting five to two, a gap of 1.7 points and the same size as the platform effect quantified above; the remaining 0.4 points is 1bk2A, which the earlier loader dropped and which this study scores as a miss. The six-configuration recomputation given in Results applies the same best-single and round-robin strategies to the 179.

### A note on comparability

These are pocket-level quantities. Much of the literature reports residue-level AUC, which asks a different question and is not comparable to any number here.

## Data and code availability

Per-candidate residue sets, centroids and overlaps for all six configurations, the collection harness, the analysis that produces every number above, and the figure generation code are at <https://github.com/mooreneural/lacuna>, archived on Zenodo under the concept DOI <https://doi.org/10.5281/zenodo.20533638>, which always resolves to the newest archived version. The released data allow coverage, conversion, top-*k* at any *k*, consensus and candidate-burden analyses to be recomputed for all six configurations without re-running any method.

The two harness defects reported here, their fixes, the machine, the commands, the per-structure regression and the invariance check on single-chain structures are recorded in benchmarks/PROVENANCE_cryptobench_full_fold_2026-11-16.md in that repository. The full-fold re-measurement quoted in Results is version 1.5.0 of the software, archived at <https://doi.org/10.5281/zenodo.24239150> [28], and its per-structure records, carrying both residue conventions, are benchmarks/cryptobench_results_chainaware.json. Benchmark outputs are released under CC-BY; the software is MIT.

## Author contributions

C.W.M. conceived the study, wrote the analysis and benchmarking code, performed all experiments, and wrote the manuscript.

