## Supplementary Tables 1 to 3 for "Cryptic binding sites are detected but not ranked: coverage, conversion, and the limits of detector consensus"

Supplementary Table 1. Greedy union of candidate sets across the six configurations, on the 179 single-chain structures analysed here, a subset of the 222 entries of the designated CryptoBench test fold (Methods). Each row adds whichever remaining configuration contributes most new coverage.

| Configurations | Union coverage | Gain |
| --- | --- | --- |
| fpocket | 73.7% |  |
| + IF-SitePred | 89.9% | +16.2 |
| + P2Rank | 91.6% | +1.7 |
| + Lacuna | 92.2% | +0.6 |
| + MDpocket | 92.2% | +0.0 |
| + Lacuna-PLM | 92.2% | +0.0 |

Supplementary Table 2. Union coverage and best single-configuration top-five under a floor on annotated site size, on the same 179 single-chain structures, a subset of the 222 entries of the designated CryptoBench test fold (Methods).

| Minimum site size | n | Union coverage | Best single top-5 |
| --- | --- | --- | --- |
| any | 179 | 92.2% | 64.2% |
| >= 6 residues | 154 | 96.8% | 73.4% |
| >= 8 residues | 138 | 98.6% | 79.7% |
| >= 10 residues | 126 | 99.2% | 83.3% |

Supplementary Table 3. Recovery under a fixed candidate budget, spent within the best single detector or spread round-robin across all four. Computed on the earlier four-detector candidate collection, fpocket, P2Rank, IF-SitePred and Lacuna under its PLM-assisted ranker, on the 178 CryptoBench test-fold targets all four ran on, and not on the 179 of Supplementary Tables 1 and 2 (Methods).

| Budget | Best single detector | Round-robin across detectors | Gain |
| --- | --- | --- | --- |
| 5 | 66.3% | 66.3% | +0.0 |
| 10 | 69.1% | 77.0% | +7.9 |
| 20 | 71.9% | 83.7% | +11.8 |
